# Lipid nanoparticle supplementation enhances host metabolism in a model symbiotic cnidarian

**DOI:** 10.1101/2025.10.06.680586

**Authors:** Ester Laurent, Jennifer L. Matthews, Lénaïc Chagnat, Chiara de Jong, Lifeng Peng, Phillip A. Cleves, David J. Suggett, Clinton A. Oakley

## Abstract

Stable cnidarian–dinoflagellate symbiosis provides the trophic foundation of coral reef ecosystems. Understanding nutrient exchange underpinning this symbiosis grows increasingly urgent as reefs face accelerating threats from climate change, and the need for time-critical interventions to improve coral health via aquaculture. Lipid nanoparticles (LNPs), widely used as delivery vehicles in biomedical science, are emerging as a promising tool to supplement coral nutrition. However, physiological impacts of LNPs on cnidarians, including uptake, nutritional value, and holobiont response, remain largely unexplored. Here, we delivered empty, phosphatidylcholine LNPs to both symbiotic and aposymbiotic *Exaiptasia diaphana*, an anemone model for corals, and analyzed the host proteomic response via mass spectrometry. LNP supplementation elicited broad proteome shifts, with notable overlap between symbiosis– and LNP-induced protein expression. Proteins involved in lipid catabolism, lipid transport, β-oxidation, lysosomal function, and protein translation were significantly more abundant, consistent with enhanced lipid processing and metabolic activity. LNP supplementation, like symbiosis, suppressed both asexual reproduction and the expression of a suite of predation– and digestion-associated venom proteins and proteases, suggesting a conserved “sated” phenotype in response to lipid supply. Variations in feeding frequency with *Artemia* had minimal impact, indicating that LNPs can be a robust supplement irrespective of primary feeding regime. These data demonstrate that adult *Exaiptasia* are capable of direct uptake of LNPs, offering a tool for probing lipid metabolism, signaling and symbiotic function in cnidarians. Moreover, the ability to manipulate host physiology using defined lipid formulations holds significant potential for advancing coral aquaculture stress resilience, including reef restoration strategies.

## Introduction

Nutrient availability plays a fundamental role in determining coral health and fitness, influencing a range of physiological and ecological traits from growth and reproduction to stress resilience (Matthews et al. 2020; Rädecker et al. 2023). In corals, nutrient exchange between the animal host and its intracellular dinoflagellate symbionts (family Symbiodiniaceae) underpins the stability and productivity of the symbiosis, which in turn supports the diverse and productive ecosystems of coral reefs (Davy et al. 2012). Cycling of nitrogen, inorganic carbon, phosphorus, and macronutrients such as carbohydrates and lipids are critical for maintaining holobiont function (Rädecker et al. 2023). Notably, Symbiodiniaceae translocate a substantial portion of photosynthetically fixed carbon to their coral hosts, much of it in the form of lipids and carbohydrates, which fuel host metabolism, tissue growth, and reproduction (Matthews et al. 2017; Hillyer et al. 2017a). In times of stress or symbiont loss (e.g., bleaching), lipid reserves become especially important, providing an energy buffer that can influence survival outcomes (Hillyer et al. 2017a, b). Understanding how nutrient availability regulates coral health is therefore critical for both advancing our understanding of coral physiology for developing effective interventions in support of reef restoration.

Exogenous feeding has been shown to enhance coral tissue biomass and growth rates in nature (Petersen et al. 2008; Conlan et al. 2017; Huffmyer et al. 2021; Rodd et al. 2022; Grottoli et al. 2025), and in aquaculture, custom designed exogenous food has been used to enhance specific coral traits such as color (Balling et al. 2008). Recently, manganese supplementation has been shown to promote photosynthetic efficiency (Biscéré et al. 2018; Moreira et al. 2025), and lipid supplementation to support coral larval swimming, settlement, and juvenile thermal tolerance (Matthews et al. 2025). Such findings suggest that supplementing nutritional inputs can improve coral performance under both ambient and stressful conditions.

Emerging technologies offer new avenues to deliver nutrient supplementation more precisely and efficiently. Lipid nanoparticles (LNPs), established in medical sciences as vehicles for RNA and drug delivery, offer a promising tool to deliver nutrients directly to cnidarian tissues (Roger et al. 2023). In addition, LNPs themselves can serve as a source for bioavailable lipids, particularly those made from phosphatidylcholine. Phosphatidylcholine is an essential structural component of eukaryotic membranes (Garrett et al. 2013) and can be broken down by phospholipases into fatty acids and subsequently used in β-oxidation and energy production (Sikorskaya 2023). Phosphatidylcholine can also be combined with glycerol to make glycerophosphocholine, an important metabolite in membrane synthesis and osmoregulation, and can serve as a precursor for lysophospholipids and other lipid signaling molecules (Rosset et al. 2021).

Beyond their potential to optimize nutrient delivery and reduce the costs of traditional feeding regimes, LNPs may also help illuminate fundamental aspects of coral metabolism and nutrient regulation (Roger et al. 2023), especially when used in controlled experiments to isolate the effects of lipid intake from those conferred by symbiosis. A major challenge in studying coral physiology is disentangling the direct effects of symbiosis from indirect effects from the nutritional contribution of the algal symbionts (Lehnert et al. 2014; Sproles et al. 2019; Rosset et al. 2021; Matthews et al. 2022; Rädecker et al. 2023). For example, changes in host immunity or gene expression during symbiosis may be due to the presence of symbionts themselves, or simply to the increased nutrient input they provide (Lehnert et al. 2014; Matthews et al. 2017). Lipid supplementation, delivered independently of symbiont activity, offers a potential tool to address this confounding factor.

In the current study, we hypothesized that adult *Exaiptasia diaphana* (Grajales and Rodríguez 2014), a model anemone for coral symbiosis research (Paxton et al. 2013), are capable of directly absorbing lipid nanoparticles. We further predicted that LNP supplementation would enhance anemone size, growth rates, and asexual reproduction, and induce proteomic shifts consistent with increased lipid metabolism. By delivering LNPs to both symbiotic and aposymbiotic *E. diaphana* and profiling host proteomic responses, we aimed to characterize the physiological impacts of exogenous lipid supplementation and hence evaluate the broader potential of lipid nanoparticles in coral reef science and management, including restoration.

## Materials and Methods

### Exaiptasia *culture*

A clonal population of *Exaiptasia diaphana*, culture identifier “NZ1”, was originally isolated from Pacific-sourced live rock and has been maintained in the laboratory at 25 °C for approximately 15 years. These anemones host the dinoflagellate *Breviolum minutum* (LaJeunesse et al. 2022; Mashini et al. 2024). Anemones were rendered aposymbiotic by repeated exposure to menthol (Matthews et al. 2016) and a stock population maintained in an aposymbiotic state for several years. All aposymbiotic anemones selected for the experiment were confirmed to be aposymbiotic by fluorescence microscopy.

Individual anemones were transferred to covered six-well plates, one anemone per well, and maintained in a climate chamber at 25 °C and 100 µmol photons m^−2^ s^−1^. Anemones were kept in 0.22 µm filtered seawater, 35 ppt, sourced from the Cook Strait. The seawater was changed and the wells cleaned at least every two days to mitigate algal growth. All experiments were performed on small (0.7–2.65 mm pedal disk) polyps. Anemones were allowed to settle in these wells for 72 h before experimentation.

### Experimental design

Symbiotic and aposymbiotic anemones were randomly assigned to experimental conditions in a two-factor design of LNP supplementation and feeding regime (Fig. 1, n = 5 per condition). LNP-supplemented anemones were supplied with 0.1 mL LNPs for one hour, three times per day, three days per week, for four weeks, with the seawater changed completely and the wells gently cleaned with a cotton swab between each addition. The seawater of control wells was also changed on the same schedule. We further used two different feeding regimes to test whether heterotrophy was necessary to balance the increased lipid availability for maximum growth, with the understanding that higher feeding rates can enhance growth (Chomsky et al. 2004). Anemones in the “low” and “high” feeding regimes were fed with freshly hatched *Artemia* nauplii once per week and twice per week, respectively, and the seawater changed the following day to minimize waste nutrient concentrations.

**Fig 1.**
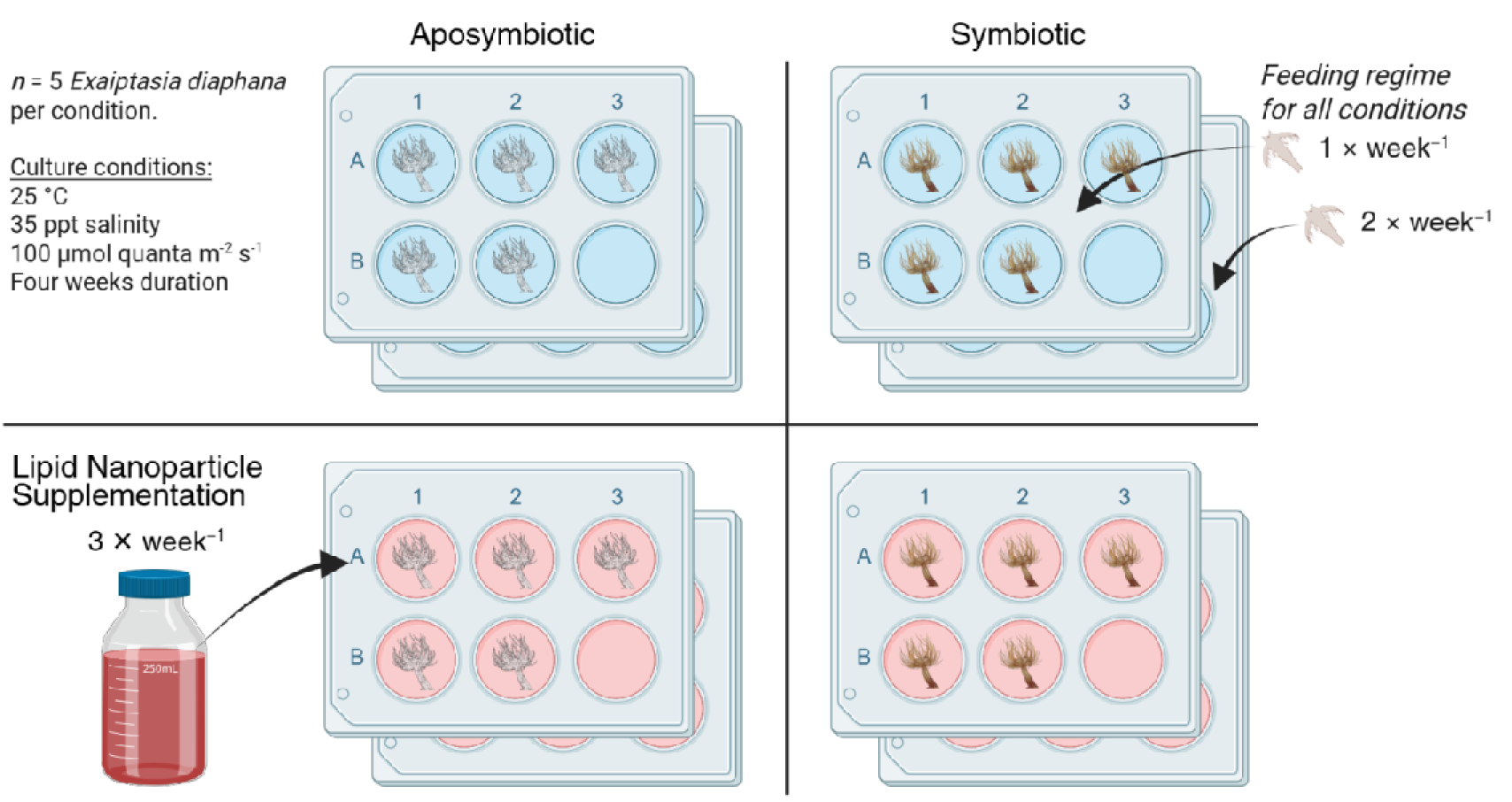
E*x*aiptasia *diaphana* were rendered aposymbiotic or maintained with their normal dinoflagellate symbiont (“Symbiotic”). Anemones were maintained individually in six-well plates. Lipid nanoparticles were supplied 3 × per week. The anemones were split between two feeding regimes, one of which received *Artemia* nauplii once and the other twice per week. Created in BioRender.

Anemone size was assessed by measurement of the pedal disk diameter with a digital caliper with 100 µm resolution, n = 18 per condition. Anemones were allowed 72 h for settlement acclimation before the first measurement, and the average of two perpendicular measurements is reported. Pedal lacerates were counted weekly by low-magnification and the lacerates then removed. The maximum quantum efficiency of photosystem II (calculated as maximal variable fluorescence / maximal fluorescence, *F_v_/F_m_* = [(*F_m_* – *F_0_*)*/F_m_*]) was assessed by pulse-amplitude modulated fluorometry (Diving-PAM, Walz, Germany), with measurements taken weekly following a 20 min dark acclimation period. Total protein mass per anemone was determined after homogenization and centrifugation (see below) by analyzing the host fraction by a fluorometric dye assay kit (Qubit, ThermoFisher Scientific). To calculate algal cell densities, the number of algae cells in the algal pellet fraction was determined by resuspending each pellet in filtered seawater, passing through a syringe needle for dispersal, and imaging using an InCell plate-reading fluorescence microscope (Cytiva). Algal cells were detected using a far-red laser for chlorophyll autofluorescence and counted using the “analyze particle” function in Fiji (v1.54).

Physiological data analysis was performed in R (Team 2021) (v 4.4.1). Data distributions were assessed using the Shapiro-Wilk test. For data following a normal distribution, analyses were conducted using ANOVA followed by Tukey’s HSD post-hoc test with Bonferroni’s correction. If the data did not follow a normal distribution, a nonparametric Kruskal-Wallis test was performed (> 2-group comparisons), followed by Dunn’s post-hoc pairwise test with Bonferroni’s correction for multiple comparisons.

### Lipid nanoparticles

The LNP solution was composed of red krill oil (Real Vitamins Ltd, Auckland, New Zealand) emulsified by soy lecithin granules (Healthy Way, Australia), following a protocol adapted from Matthews *et al*. (2025a). One gram lecithin was dissolved in 250 mL water at 55 °C, followed by the gradual addition of 2.5 g total lipid (10 mg mL^−1^ final), while continuously stirring. Tween-20 was then added gradually to 2% total by volume. After 30 min the mixture was stored at 4 °C, and warmed and agitated before use. By the manufacturer’s specifications, the krill oil was composed of 30% omega-3 fatty acids by weight, including eicosapentaenoic acid (15% total) and docosahexaenoic acid (8%), as well as phospholipids (41.3%) and astaxanthin (0.13%). The zeta potential of the LNP suspension was −25.07, indicating nanoparticle stability, with an average particle size of 463.1 nm upon measurement using a Malvern Zetasizer Nano ZS instrument (Malvern Panalytical, UK) featuring backscattering detection at a consistent 173° scattering angle. The apparatus was equipped with a He–Ne laser (wavelength = 632.8 nm) and maintained at a temperature of 25°C. LNPs were measured using Zetasizer folded capillary cells DTS 1060 (Malvern Panalytical, UK). Data analysis was processed using the instrumental Malvern’s DTS software to obtain the mean ZP value. All LNP measurements were performed in triplicate.

### Proteomic sample preparation

*Exaiptasia* were processed similarly to Oakley *et al*. (2023). After four weeks, anemones were removed from the plates into microcentrifuge tubes. Salts were removed by washing with 1 mL 4°C HPLC-grade water, resuspended in 500 uL HPLC-grade water, and mechanically and rapidly homogenized with approximately 10 passes with a Dounce homogenizer to lyse host cells. The symbiont fraction was separated by centrifugation (300 *g* × 30 s) and the supernatant, containing host protein, was transferred to a new tube. The algal pellet was frozen at −80 °C until used for symbiont cell density measurements. Host protein was dissolved and denatured by the addition of sodium deoxycholate to a final concentration of 5% w/w, in 100 mM triethylammonium bicarbonate buffer (TEAB, pH 8.5). β-mercaptoethanol was added to 20 µM final concentration as a reductant to further denature and solubilize proteins. Samples were further denatured at 90−95 °C for 20 min and then disrupted using an ultrasonicator probe (20 × 2 s pulses). The sample was transferred to a 0.5 mL 30 kDa molecular weight cutoff filter (Amicon Ultra, Merck Millepore) for filter-aided sample preparation (Wiśniewski et al. 2009).

The protein sample was washed twice by centrifugation and resuspended with 380 µL 100 mM TEAB in the filter (14,000 *g* × 20 min), before resuspension in 400 µL 100 mM TEAB. A 10 µL subsample was taken, acidified to 1% formic acid by volume, the deoxycholate precipitate removed by centrifugation (10,000 *g* × 2 min), and the protein in the supernatant quantified (Qubit 2.0, ThermoFisher Scientific, USA). 100 µg protein from the remaining sample was reduced with 10 mM β-mercaptoethanol (final) at 37 °C for 10 min, then alkylated with 20 mM acrylamide (final) at room temperature for 20 min. Protein was then digested with 2 µg trypsin (Pierce) overnight at 37 °C in the filter. Peptides were separated from undissolved protein by centrifugation (14,000 *g* × 20 min), acidified with 1% formic acid (final) and centrifuged (10,000 *g* × 2 min) to remove deoxycholate. Peptides were then desalted with C18 tips (Omix Bond Elut, Agilent Technologies), dried by vacuum centrifugation, and stored at 4 °C until analysis.

### Mass spectrometry

Peptides were resuspended in 0.1% formic acid, dissolved for 30 min at 37 °C, and analyzed by liquid chromatography-tandem mass spectrometry. 250 ng total peptide were loaded onto a 50 cm C18 column (Acclaim PepMap C18 #164570, ThermoFisher Scientific, 3 μm particle size, 100 Å) and Ultimate 3000 liquid chromatograph (ThermoFisher Scientific). Peptides were eluted by a 75 min gradient from 5%–35% buffer B (buffer A: 0.1% formic acid; buffer B: 80% acetonitrile, 0.1% formic acid) at 300 nL min^−1^ and 55 °C. Peptides were analyzed with a Lumos Tribrid Hybrid mass spectrometer (ThermoFisher Scientific) by electrospray ionization at a 1.8 kV spray voltage and a resolution of 120,000. The top 20 MS peaks over a scan range of 375–1400 m/z were analyzed by the orbitrap, rejecting +1 charge states and with dynamic exclusion enabled (60 s). Peptides were fragmented by collision-induced dissociation and fragments analyzed by the ion trap. Instruments were operated with Chromeleon (v7.3.1), Xcalibur (v4.7), and Tune (v4.1.4244, ThermoFisher Scientific).

Protein identification was conducted using the MSFragger search engine (v4.1) in FragPipe (v22.0) against protein sequences derived from the *Exaiptasia diaphana* genome (Baumgarten et al. 2015) (v1.1, NCBI accession PRJNA261862). False discovery rate (FDR) thresholds were set at 1% for peptide and protein search matches, and a minimum of two peptides per protein were required for identification. Searches assumed trypsin digestion with a maximum of two missed cleavages. Oxidation of methionine and acetylation of the protein N-terminus were specified as variable modifications, and propionamidylation of cysteine was specified as a fixed modification. Proteins were quantified using the MaxLFQ method in IonQuant (v1.10.27) in FragPipe with ‘match between runs’ enabled and intensities normalized across runs.

### Protein abundance analysis

Comparison of protein abundances between treatments was performed similarly to Mashini *et al*. (2023). Known false matches/decoys and typical contaminants were removed and label-free quantification intensity values were log_2_-transformed. Differentially abundant proteins (DAPs) were then identified using limma (Ritchie et al. 2015), with an FDR of *q* < 0.05. Principal component analysis was performed in ClustVis (Metsalu and Vilo 2015), and area-proportional Venn diagrams generated using the eulerr package in R (Larsson et al. 2024).

### Gene set enrichment analysis

The biological functions of DAPs were further explored by gene set enrichment analysis using topGO (Alexa et al. 2006; Alexa and Rahnenfuhrer 2024). For each comparison, the FDR values generated by limma of all detected proteins were used as the significant gene list. Significant gene ontology (GO) terms were determined by the weight01 algorithm and the Kolmogorov-Smirnov test statistic (“ks”), with a node size of 5 and a FDR requirement (“geneSelectionFun”) of 0.05. Significant GO terms were those with a p-value < 0.05. When analyzing the set of DAPs which responded in the same direction to both symbiosis and LNP supplementation, the FDR of all DAPs in this set was manually set to 0.01, the FDR of all other proteins set to 1.0, and the Fisher test statistic (“fisher”) used.

## Results

### Exaiptasia physiology with LNP supplementation

Total protein mass per anemone did not increase with addition of LNP (supplementation) or presence of Symbiodiniaceae (symbiotic state; Fig. 2a,b); however, pedal disks were smaller under both states compared to LNP-free or aposymbiotic (Fig. 2c). Pedal disk size was reduced further in symbiosis by LNP supplementation (Fig. 2d). Total protein mass per anemone and pedal disk size were greater amongst anemones fed twice per week. Combined LNP supplementation and high feeding increased total protein mass in aposymbiotic anemones when compared to symbiotic anemones with low feeding and no LNP supplementation (Supp. Fig. 1a). We observed no mortality during the experiment. Pedal lacerate generation was significantly reduced by both symbiosis and LNP supplementation (Fig. 2e, f), with the number of pedal lacerates generated by LNP-supplemented aposymbiotic anemones not significantly different from symbiotic anemones (Fig. 2f). Pedal lacerate generation was greater amongst anemones fed twice per week (Supp. Fig. 1c).

**Fig 2.**
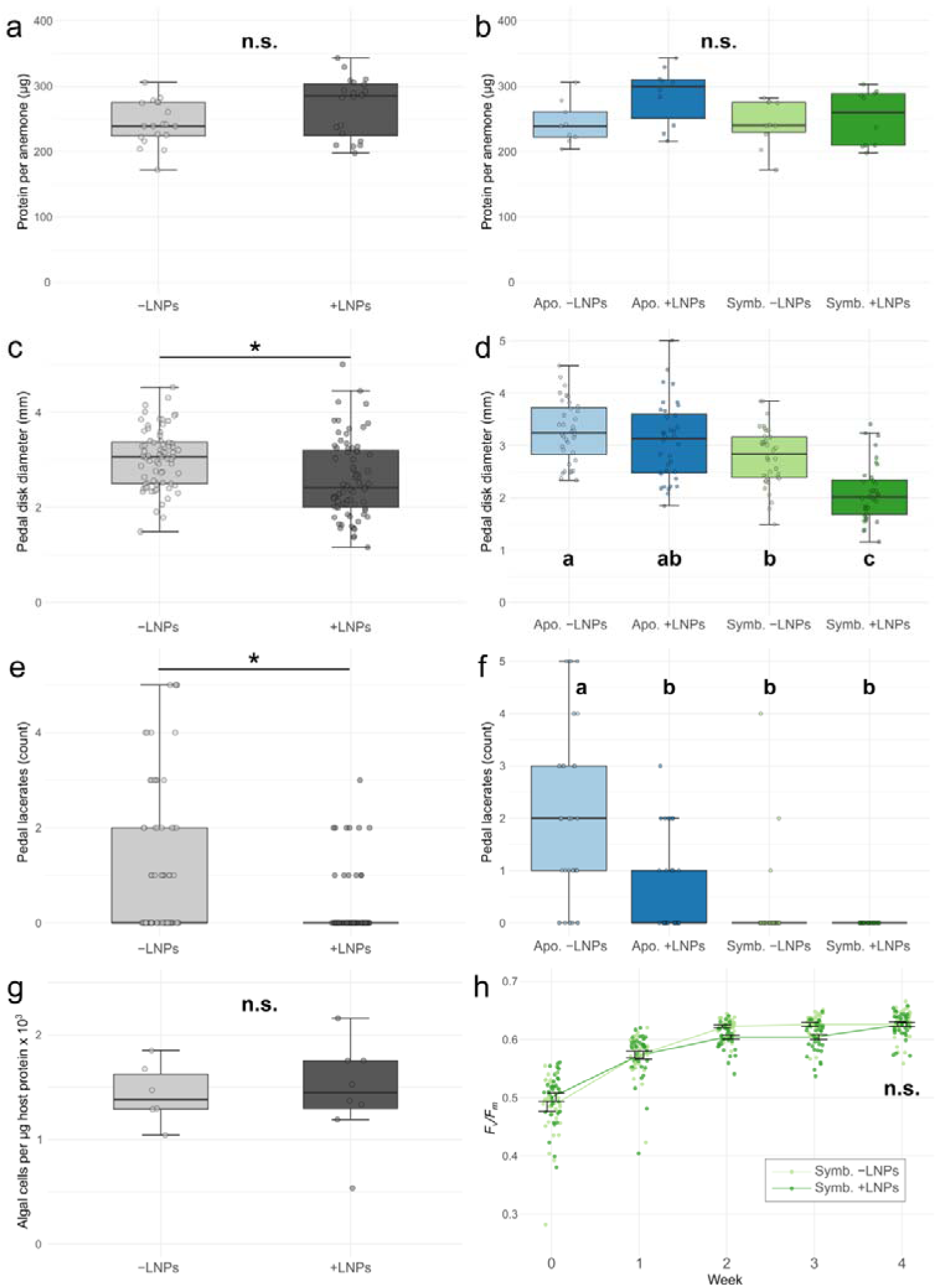
Physiological responses of *Exaiptasia diaphana* to lipid nanoparticle (LNP) supplementation. a, b: Protein mass per anemone at the end of the four-week experiment. c, d: Pedal disk size at the end of the four-week experiment. e, f: Total counts of pedal lacerates across the four-week experiment. g: Cell density of dinoflagellate symbionts, normalized to host protein. Feeding regimes have been combined. h: *F_v_/F_m_* of symbiotic anemones across the four-week experiment (*n* = 18). Conditions with asterisks (one-way ANOVA, *p* < 0.05) or different letters (two-way ANOVA with Tukey multiple comparison, *p* < 0.05) are significantly different. Apo.: aposymbiotic; Symb.: *Exaiptasia* hosting *Breviolum minutum*

Symbiotic, LNP-supplemented anemones did not produce any pedal lacerates throughout the experiment. Algal symbiont cell densities did not respond to LNP supplementation (Fig. 2g). LNP supplementation resulted in minimal differences in the symbiont maximum quantum efficiency of photosystem II (*F_v_/F_m_*), after and throughout an initial period of photoacclimation to greater irradiance (Fig. 2h).

### Proteome results

Of the 4,357 identified *Exaiptasia* proteins, the effect of symbiosis (1,622 DAPs, Fig. 3b) was larger than that of LNP supplementation (406 DAPs). The effects of LNP supplementation and symbiosis on the *Exaiptasia* proteome are visualized by principal component analysis (Fig. 3a) indicating a consistent effect of LNP supplementation on both symbiotic and aposymbiotic proteomes. Feeding regime had a negligible effect on the proteome (1 DAP), which was consistent when comparing low and high feeding regimes within each symbiotic state or LNP supplementation treatment (Table 1). Based on these minimal effects of feeding regime on both the proteome and observed physiology, we have combined sample data from the low and high feeding regimes within each condition for the remaining analyses. Full protein abundance data and comparisons between treatments are available as Supp. Tables 1–3, and spectral data are available at the ProteomeXchange Consortium, identifier PXD066136.

**Fig 3.**
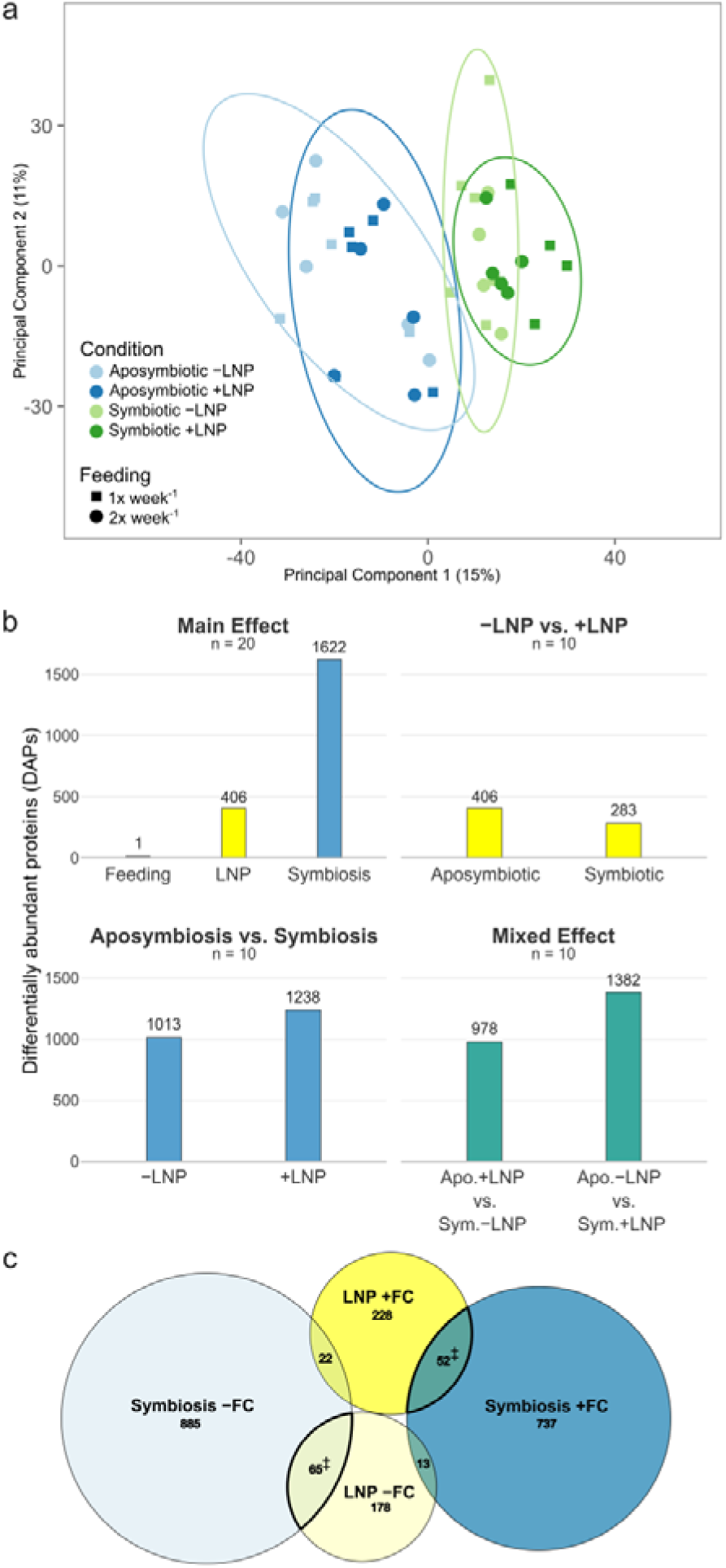
a: Principal component analysis of protein abundances of aposymbiotic and symbiotic *Exaiptasia diaphana*, with or without lipid nanoparticle (LNP) supplementation. Each point represents one biological sample. Prediction ellipses represent 95% confidence intervals. Symbiotic state and LNP supplementation conditions are noted by color, while *Artemia* feeding regime is indicated by point shape. b: Proteins which were significantly differentially abundant (FDR, *q* < 0.05) between conditions when considering either all samples (“main effect”, n = 20) or subsets (n = 10). c) Proteins which were significantly differentially abundant due to symbiosis or LNP supplementation, presented by direction of fold-change. n = 20 for each comparison. Symbiosis –FC/+FC: Proteins which were less/more abundant, respectively, in symbiotic *Exaiptasia* when compared to aposymbiotic anemones. LNP – FC/+FC: Proteins which were less/more abundant, respectively, in anemones supplemented with LNPs. ‡Protein sets which were selected for further gene ontology enrichment analysis (see Figs. 4 and 6). See Supp. Table 9 for full protein lists

**Table 1.** Significant differentially abundant (FDR, *q* < 0.05) proteins of *Exaiptasia diaphana* between symbiosis, lipid nanoparticle (LNP) supplementation, and feeding conditions. Apo.: Aposymbiotic; Symb.; Symbiotic. LF: Low Feeding (1× week^-1^), HF: High Feeding (2× week^-1^).

|  | Apo.<br>LF<br>-LNP | Apo.<br>HF<br>-LNP | Apo.<br>LF<br>+LNP | Apo.<br>HF<br>+LNP | Symb.<br>LF<br>-LNP | Symb.<br>HF<br>-LNP | Symb.<br>LF<br>+LNP |
| --- | --- | --- | --- | --- | --- | --- | --- |
| Apo. LF -LNP | - |  |  |  |  |  |  |
| Apo. HF -LNP | 0 | - |  |  |  |  |  |
| Apo. LF +LNP | 137 | 146 | - |  |  |  |  |
| Apo. HF +LNP | 177 | 93 | 0 | - |  |  |  |
| Symb. LF -LNP | 360 | 287 | 423 | 409 | - |  |  |
| Symb. HF -LNP | 614 | 295 | 463 | 296 | 1 | - |  |
| Symb. LF +LNP | 873 | 758 | 532 | 598 | 155 | 183 | - |
| Symb. HF +LNP | 653 | 497 | 277 | 250 | 113 | 78 | 2 |

### Proteome effects of LNP supplementation

The 406 proteins that responded to LNP supplementation were significantly enriched (*p* < 0.05, topGO) in protein translation, proteolysis, toxin activity, immune response, lipid catabolism, and apoptosis regulation processes (Figs. 3–5). Cell components enriched in these proteins included the lysosome, nematocyst, proteasome complex and mitochondrion (Fig. 4 and 5). DAPs involved in protein translation, immune response, lipid catabolism, and apoptotic regulation were largely more abundant in response to LNP supplementation, while venoms and proteases were less abundant (Fig. 5).

**Fig 4.**
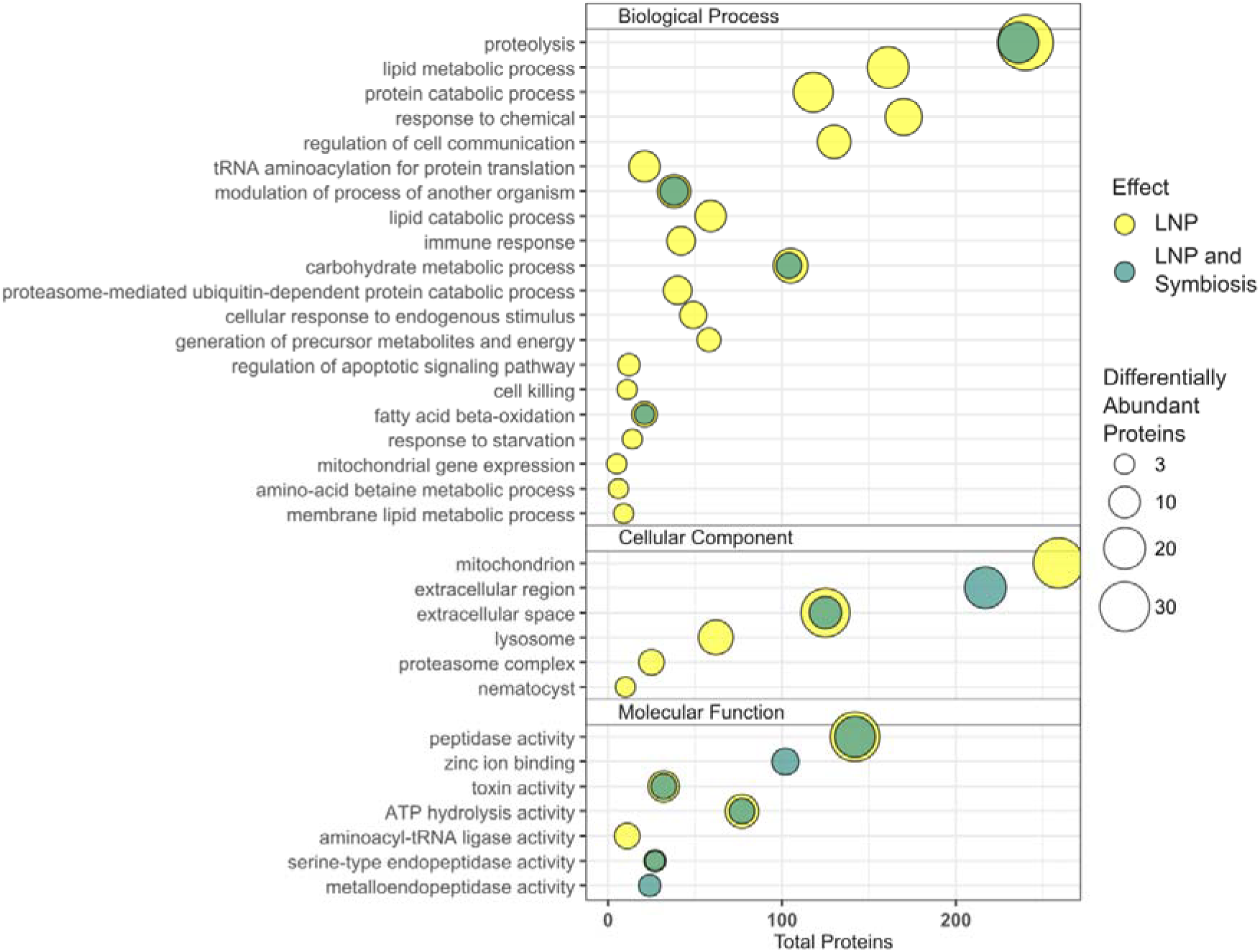
Selected gene ontology (GO) terms which were significantly enriched (*p* < 0.05) among proteins which were differentially abundant (*q* < 0.05) in response to lipid nanoparticle (LNP) supplementation (yellow) and among proteins which were more abundant in response to both LNP supplementation and symbiosis (green). Terms are arranged by GO domain and total number of detected proteins annotated with each term. Terms with less than three differentially abundant proteins have been omitted. Full results are available in Supplementary Tables 4–6

**Fig 5.**
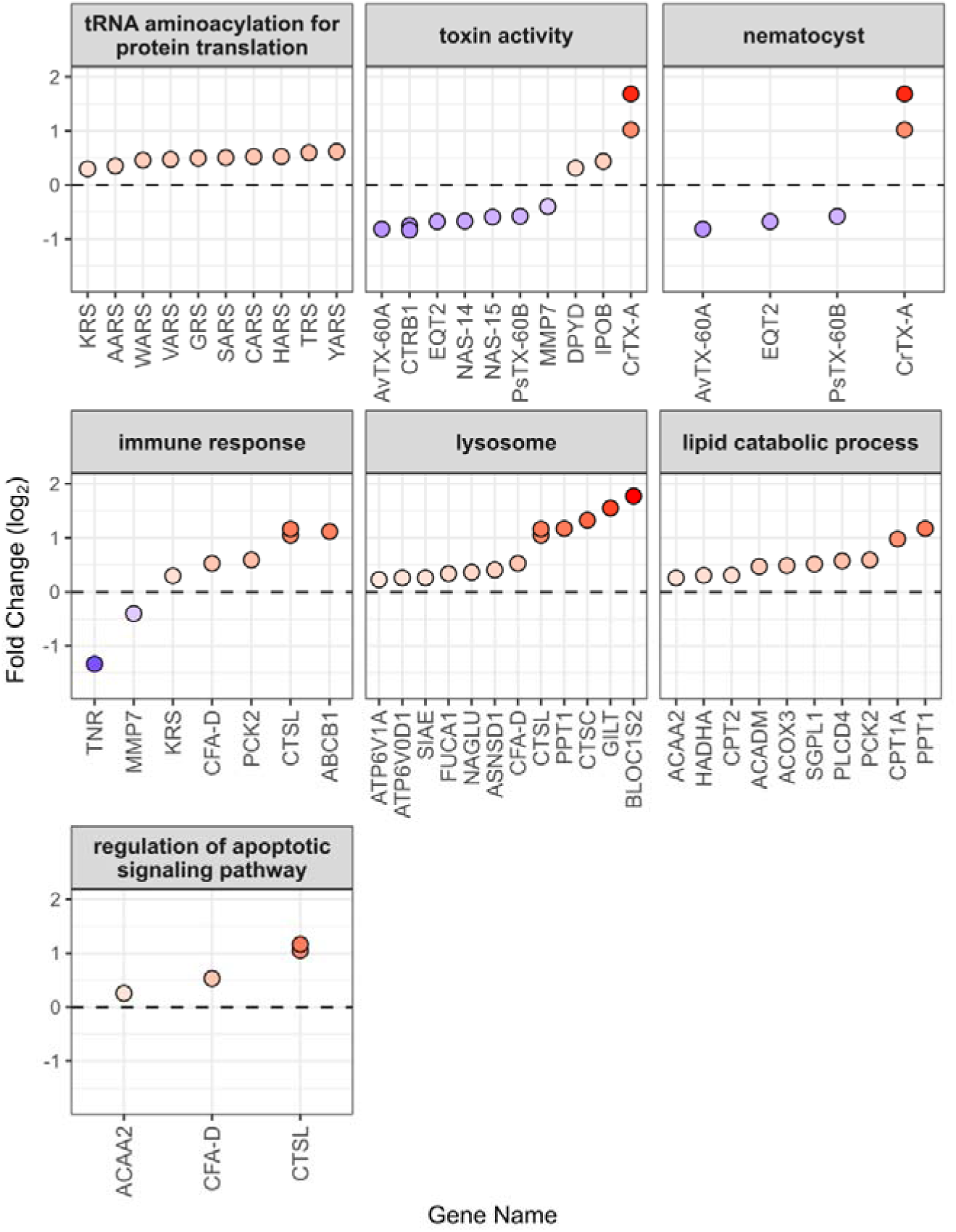
Changes in *Exaiptasia diaphana* protein abundance and significantly enriched gene ontology terms in response to LNP supplementation. Proteins shown are differentially abundant (FDR, *q* < 0.05), with positive values indicating increased abundance in response to LNP supplementation. Selected significant (*p* < 0.05) gene ontology terms are displayed. Full data are available in Supp. Table 7

### Proteome effects of symbiosis

The symbiotic state of *Exaiptasia* had a large effect on the proteome (1,622 DAPS of 4,357 total identified proteins, 37.2%), as previously described (Oakley et al. 2016; Sproles et al. 2019). Our findings here are consistent with these earlier works. Effect size was substantially larger for Symbiodiniaceae symbiosis than for LNP supplementation (Fig. 3b, Table 1). 133 GO terms were significantly enriched (*p* < 0.05, topGO) across all domains. Significant biological processes responding to symbiotic state include translation, chitin catabolism, innate immune response, regulation of intracellular transport, protein folding, and fatty acid β-oxidation, while significant cellular component and molecular functions include the collagen-containing extracellular matrix, ribosome, and toxin activity (Supp. Table 5).

### Similarities between LNP supplementation and symbiosis

To further assess whether LNP supplementation and symbiosis had similar or congruent effects on the host proteome, we performed further gene ontology enrichment analysis on DAPs which responded to both symbiosis and LNP supplementation (*q* < 0.05 for both) with abundance fold change in the same direction, positive or negative (Figs. 3c, 4, 6). Overlapping DAPs were enriched in fatty acyl-CoA hydrolase activity and lipid droplet terms with positive fold change. Proteolysis and toxin activity, composed of a range of proteases and peptidases, showed bidirectional shifts. Broadly, 26S proteasome components were more abundant and a suite of other proteases/peptidases less abundant during symbiosis or LNP supplementation.

**Fig 6.**
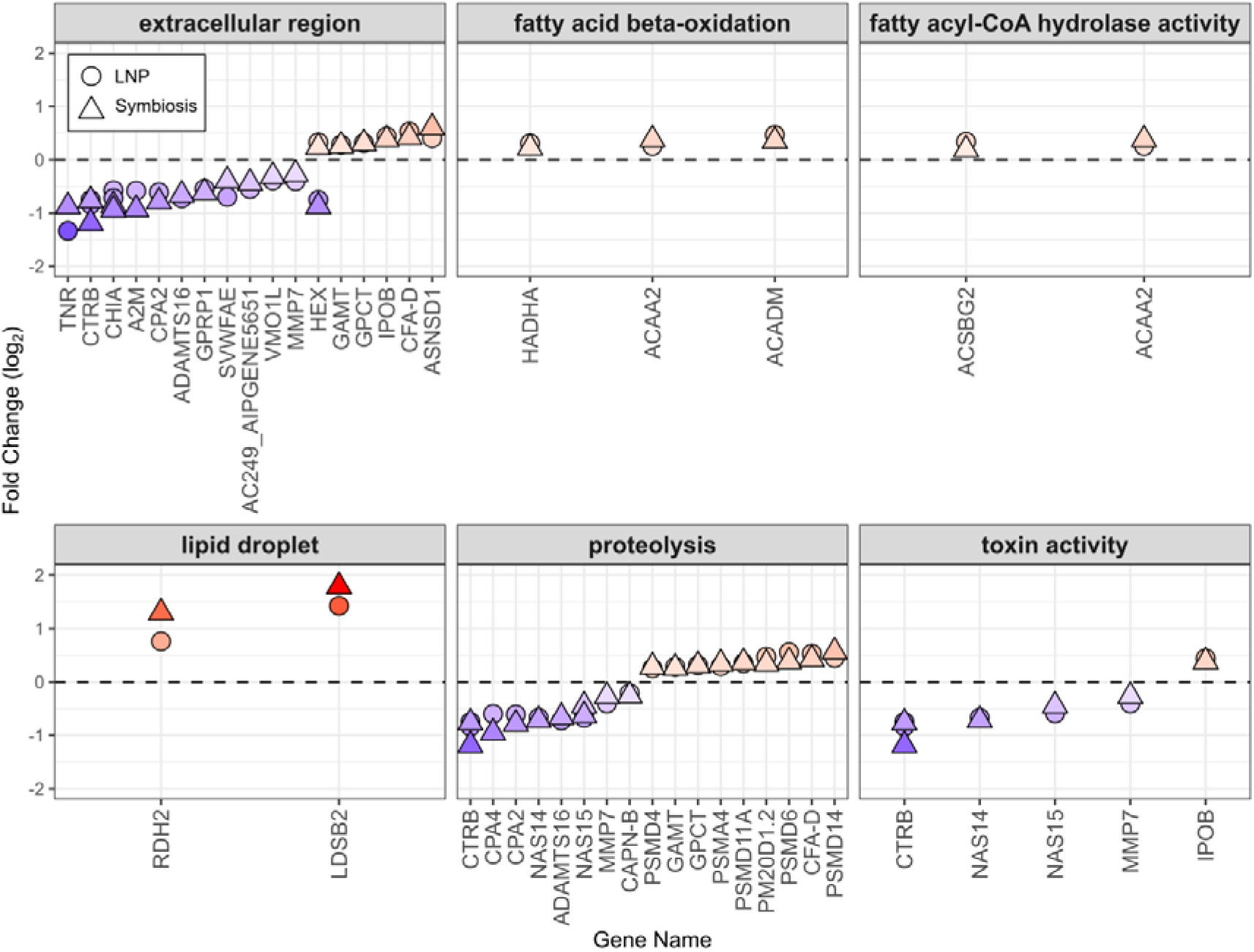
Congruent changes in *Exaiptasia diaphana* protein abundance in response to lipid nanoparticle supplementation and symbiotic state. Proteins shown are differentially abundant (FDR, *q* < 0.05) in both LNP supplementation and symbiotic conditions, with the same direction of fold change in both. Proteins are grouped by gene ontology term, all of which were significantly enriched (p < 0.05) amongst these proteins (Table 4). Positive values indicate increased abundance in response to both LNP supplementation and symbiosis. Full data are available in Supp. Table 8

## Discussion

### LNP supplementation increases cnidarian lipid catabolism, protein translation, and central metabolism

Direct LNP supplementation had clear and consistent effects on the proteome of adult *Exaiptasia diaphana*, regardless of symbiotic state. Notably, ten proteins annotated with the Gene Ontology term “lipid catabolic process” were significantly more abundant following LNP supplementation (Fig. 5), including all enzymes required for mitochondrial fatty acid β-oxidation. Central to this process are carnitine o-palmitoyltransferase 1a (CPT1a; log_2_ fold change [FC] = 0.98) and carnitine o-palmitoyltransferase 2 (CPT2, FC = 0.31), key components of the carnitine shuttle system, which facilitates the import of long-chain fatty acids into the mitochondria for β-oxidation (Longo et al. 2016). Within mitochondria, the fatty acyl-CoAs are catabolized through the β-oxidation cycle. Enzymes catalyzing all four steps of this cycle were more abundant in response to LNP supplementation: medium-chain acyl-CoA dehydrogenase (ACADM, FC = 0.47) initiates the cycle, followed by two steps conducted by the mitochondrial trifunctional enzyme (HADHA, FC = 0.30), followed by cleavage by 3-ketoacyl-CoA thiolase (ACAA2, FC = 0.26) to produce acetyl-CoA, which drives the citric acid cycle (Houten and Wanders 2010). Such concerted upregulation provides strong evidence that LNP supplementation enhances mitochondrial lipid metabolism in the host.

*Exaiptasia* proteins of the TCA cycle, which utilizes the acetyl-CoA produced by β-oxidation, also responded to LNP supplementation. Phosphoenolpyruvate carboxykinase (PCK2, FC = 0.59) catalyzes the formation of oxaloacetate, which combines with acetyl-CoA to form citrate in a reaction catalyzed by citrate synthase in the first step of the TCA cycle (Yang et al. 2009). Citrate is then converted to isocitrate by aconitate hydratase (FC = 0.41, Supp. Table 1), where it can either continue the TCA cycle via isocitrate dehydrogenase (FC = 0.27), or enter the two-step glyoxylate cycle via isocitrate lyase (FC = 1.09). The glyoxylate cycle bypasses the decarboxylation steps of the TCA cycle and enables gluoconeogenesis from fatty acid-derived acetyl-CoA – an adaptation associated with efficient carbon retention (Dunn et al. 2009). These enzymes were all more abundant in response to LNP supplementation. A lipid storage droplet surface-binding protein (FC = 1.43, Supp. Table 1) also showed increased abundance, suggesting an enhanced capacity for intracellular lipid storage. Taken together, these responses indicate that LNP supplementation not only increases lipid uptake and mitochondrial fatty acid catabolism, but also promotes metabolic processing and potentially storage. This proteomic response therefore demonstrates the capacity of *Exaiptasia* to utilize exogenous lipid sources, and provides a mechanistic framework for understanding how LNP supplementation can modulate cnidarian metabolism at the molecular level.

LNP supplementation further increased the abundance of ten amino acid–tRNA ligases (FC = 0.30–0.62, Fig. 5), suggesting an overall upregulation of translational machinery. These ligases catalyze the charging of specific tRNAs with their cognate amino acids (Park et al. 2008), a foundational step in protein synthesis. The coordinated increase in these ligases implies elevated translational potential, although we did not observe an increase in protein biomass under the current experiment duration (except for aposymbiotic anemones with high feeding, Supp. Fig. 1a).

### Lysosomal remodeling in response to LNP supplementation

Lysosomes are acidic cell compartments specialized for the efficient breakdown of proteins, carbohydrates, and lipids by a range of hydrolases (Xu and Ren 2015). Thirteen lysosomal-associated proteins increased in abundance following LNP supplementation, consistent with enhanced intracellular degradation and nutrient recycling (Fig. 5). Two V-type proton ATPase subunits (ATP6V1A and ATP6V0D1) showed modest increases (FC = 0.22 and 0.26 respectively) supporting lysosomal acidification (Fig. 5). Hydrolases involved in carbohydrate and glycoprotein metabolism (SIAE, NAGLU, FUCA1, ASNSD1, FC = 0.26– 0.40) were similarly upregulated. The proteases cathepsin L (CTSL, FC = 1.05 and 1.16) and lysosomal aspartic protease (CTSC, FC = 1.33) were also more abundant; these have previously been found to be less abundant in aposymbiotic *Exaiptasia* relative to symbiotic animals (Sproles et al. 2019). Additionally, Niemann-Pick type 2 (NPC2), which facilitates intracellular sterol transport, was more abundant following LNP supplementation (Ebner et al. 2025) (Supp. Table 1). Such responses are especially relevant given that the symbiosome membrane, which surrounds dinoflagellates in this symbiosis and mediates all nutrient exchange and signaling between partners, is a derived late endosomal membrane which resembles a lysosome (Fitt and Trench 1983; Mohamed et al. 2016; Voss et al. 2023). These results identify the lysosome as a central site of lipid catabolism in *Exaiptasia*, and may also reflect the shared properties of lysosomes, late endosomes, and the symbiosome membrane (Voss et al. 2023).

### Reduction in venoms and proteases in response to LNP supplementation

LNP supplementation, and symbiosis, also resulted in a notable decline in several venom proteins and digestive proteases. Key nematocyst-localized pore-forming toxins, including equinatoxin-2 (EQT2, −0.68 FC) and venoms PsTX-60B and AvTX-60A (−0.58 FC and −0.81 FC), were less abundant in LNP-supplemented anemones (Fig. 5). These toxins breach prey cell membranes, inducing osmotic lysis (Oshiro et al. 2004; Satoh et al. 2007; Rivera-de-Torre et al. 2019), and are integral to cnidarian predation. In contrast, the venom CrTX-A (Nagai et al. 2002) was more abundant (−1.03 and 1.69FC, Fig. 5), suggesting differential roles or selective regulation of venom in response to metabolic state.

Acontial filaments within the gastrovascular cavity are armed with nematocysts in *Exaiptasia* (Schlesinger et al. 2009), where post-capture prey digestion is further aided by proteases. We observed a simultaneous decline in several proteases associated with prey digestion, including two chymotrypsinogens (CTRB), two zinc metalloproteinases, NAS-14 and NAS-15, and matrilysin (MMP7). Coordinated reduction in both venom and digestive protease abundance suggests a shift toward a “sated” phenotype characterized by reduced dependence on predation. Nematocysts, being single-use organelles which are highly abundant in the cnidarian epidermis, are inherently costly to produce in terms of protein content (He et al. 2023). Changing venom content may be a mechanism of tuning the predation response to reflect the animal’s nutritional state. We therefore propose that anemones suffering from energy deprivation, either through starvation or loss of dinoflagellate endosymbionts, express greater concentrations of venoms and proteases, and are therefore more effective at predation and prey digestion for sustenance.

### Congruency of LNP supplementation and symbiosis

Supply of photosynthetically-derived carbohydrates, amino acids, and lipids are essential for host metabolism, growth and survival (Muscatine and Porter 1977; Rädecker et al. 2023).

Lipids in particular, have emerged as critical players not only in energy storage and membrane synthesis, but also in mediating symbiosome structures, symbiotic signaling, and metabolic regulation (Matthews et al. 2017; Voss et al. 2023; Gamba et al. 2024). We sought to compare the proteomic responses of *E. diaphana* to LNP supplementation with those induced by a stable symbiotic state. A scenario of the two treatments producing similar patterns in protein expression could enhance our understanding of how lipid trafficking supports a functional symbiosis and of liposomal processes involved in symbiosome formation and stability.

Both LNP supplementation and symbiosis resulted in greater abundance of proteins involved in lipid metabolism, transport, and storage. One of the most prominent shared responses was an increase in lipid storage droplet-binding protein 2 (LSDB2), which in *Drosophila* is localized to the surface of lipid storage droplets, the primary site of neutral lipid accumulation (Teixeira et al. 2003). Likewise, key enzymes involved in fatty acid β-oxidation (Houten and Wanders 2010), including carnitine O-acetyltransferase, both the alpha and beta subunits of the mitochondrial trifunctional enzyme (HADHA and HADHB), and 3-ketoacyl-CoA thiolase (ACAA2), were more abundant under both conditions. A very long-chain acyl-CoA dehydrogenase, essential for initiating the oxidation of extended fatty acid chains, also increased (Figs. 6, 7). HADHA, ACAA2, and acyl-CoA dehydrogenases form the core of mitochondrial lipid catabolism and their upregulation under both treatments suggests that lipids delivered via LNPs are bioavailable, readily metabolized by the host, and capable of mimicking the metabolic state of a symbiotic cnidarian. This outcome therefore highlights the potential utility of LNPs in influencing the metabolic state of corals in both experimental and applied settings (Voss et al. 2023).

Further evidence for convergence between anemones hosting dinoflagellate symbionts and aposymbiotic anemones supplemented with LNPs comes from observations of the Niemann-Pick Type C2 (NPC2, or epididymal secretory protein E1) protein family, known for their role in sterols transport and symbiosis specificity in *Exaiptasia* and other cnidarians (Dani et al. 2014; Oakley et al. 2016; Hambleton et al. 2019; Sproles et al. 2019). NPC2 proteins bind sterols and facilitate their movement across membranes, with some isoforms localized to the symbiosome and others functioning throughout the host cytoplasm (Hambleton et al. 2019).

We identified four distinct NPC2 proteins (Supp. Tables 1 and 2): three increased in symbiotic animals, including one with a 3.77-fold change (KXJ25141.1), consistent with its role in symbiosome function (Voss et al. 2023). Notably, one NPC2 isoform (KXJ13076.1) was significantly upregulated in response to LNP supplementation alone (FC = 1.67), independent of symbiosis. Differences between NPC2 responses here support the hypothesis that some NPC2s may be specific to symbiotic function, while others may participate in broader host sterol transport, likely within lysosomal or endosomal compartments (Dani et al. 2014; Hambleton et al. 2019; Ebner et al. 2025). Distinguishing the roles and localization of these NPC2 isoforms may provide valuable insight into lipid dynamics and host–symbiont compatibility.

In both LNP-supplemented and symbiotic anemones, we also observed reduced abundance of multiple proteases associated with prey capture and digestion. Multiple proteases included chymotrypsinogen (CTRB), calpain (CAPN-B), matrix metalloproteinase 7 (MMP7), and five zinc metalloproteinases (ADAMTS16, CPA2, CPA4, NAS14, NAS15, Fig. 6). We propose that these protease enzymes (in addition to their increased abundances of venoms) are upregulated in nutrient-deprived states, such as aposymbiosis or bleaching (Houlbrèque and Ferrier-Pagès 2009), to enhance heterotrophic feeding capacity. Suppression of these protease enzymes in response to LNPs or symbiont-derived nutrients further suggests a shift toward a metabolically replete or “sated” phenotype, where the energetic investment in prey digestion and nematocyst synthesis may be unnecessary.

Simultaneously, both treatments increased the abundance of several protease subunits, including four from the 26S proteasome (PSMD6, PSMD14, PSMD11A, PSMD4) and one from the 20S proteasome (PSMA4) (Fig. 6, Supp. Tables 1 and 2). Proteasomes are critical for the selective degradation of proteins by ubiquitination, regulating protein quality control, cell cycle progression, and metabolic homeostasis (Schweitzer et al. 2016). Notably, PSMD14 (RPN11), a deubiquitinating enzyme required for recycling ubiquitin from substrates targeted for degradation (de Poot et al. 2017), was elevated under both conditions, which may reflect enhanced proteome maintenance, removal of damaged proteins (VerPlank et al. 2019), or increased metabolic turnover, responses enabled by improved nutrient status. However, the specific role of proteasomal activity in this context warrants further investigation.

Collectively the various overlapping proteomic responses we observed demonstrate that LNP supplementation not only supports host lipid metabolism and storage, but also induces shifts in host physiology more like that of a successful cnidarian–dinoflagellate symbiosis. The findings therefore suggest that LNPs can serve as both a practical intervention to support cnidarian health in aquaculture and reef restoration, and a research tool to decouple nutritional status from symbiotic state.

### Utility of lipid nanoparticles in coral biology, aquaculture, and bleaching recovery

Our current study provides the first evidence that adult cnidarians, even in the absence of symbionts, can absorb and metabolize LNPs delivered directly into seawater (Fig. 7). These findings advance our understanding of cnidarian lipid biology and offer a promising new avenue for both experimental and applied use. Notably, the LNPs used here were composed solely of phosphatidylcholine and lacked cargo to induce specific biological effects, such as proteins or plasmids, but were still sufficient to induce broad proteomic shifts across metabolism, digestion, and protein turnover, strongly indicating that empty LNPs themselves are a bioavailable and physiologically impactful nutrient source.

**Fig 7.**
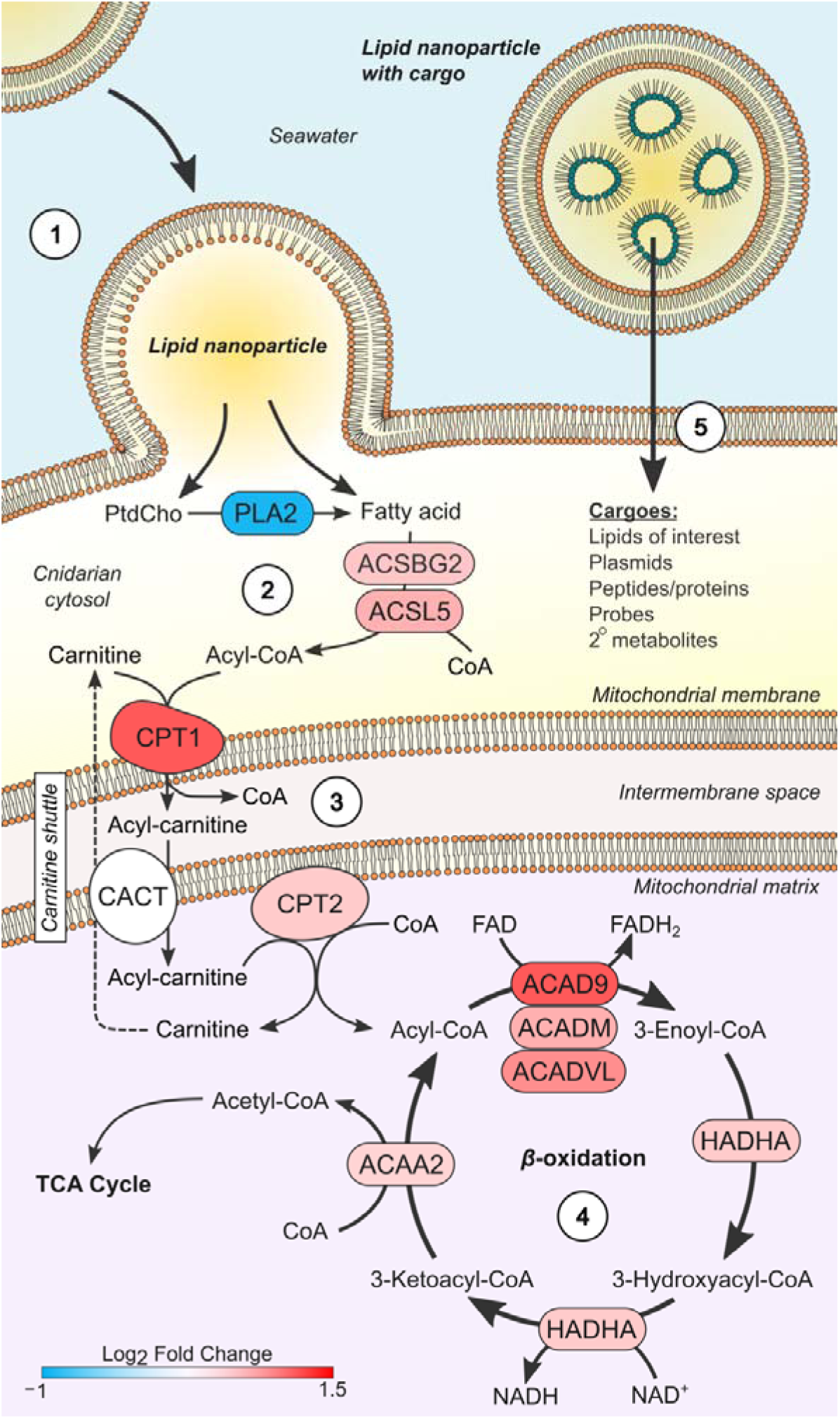
Model of lipid nanoparticle absorption by adult *Exaiptasia diaphana* and their effect on lipid metabolism. 1) Lipid nanoparticles suspended in seawater are directly absorbed by the host epidermis. 2) Phosphatidylcholine is cleaved to release fatty acids, which, with the fatty acids directly incorporated into the nanoparticle, are activated by fatty acid-CoA ligases to acyl-CoA. 3) Acyl-CoA is converted to acyl-carnitine and transported into the mitochondrial matrix by the carnitine shuttle system, where it is converted back to acyl-CoA. 4) Acyl-CoA enters β-oxidation to generate acetyl-CoA, which enters the citric acid/TCA cycle. 5) Other potential cargoes may be considered to support cnidarian symbiosis and conservation research. Changes in protein abundance due to lipid nanoparticle supplementation are indicated by color, where red indicates greater abundance. PtdCho: phosphatidylcholine, CoA: coenzyme A. Full data, including protein names, are available in Supp. Table 10

These proteomic, phenotypic, and behavioral changes in *Exaiptasia* elevates LNPs from inert delivery vehicles to functional dietary supplements, capable of altering the physiological state of the host. The observed convergence between LNP supplementation and the metabolic phenotype of symbiotic animals, including upregulation of lipid metabolism pathways, suppression of predation-related proteases, and increased proteasome activity, suggests that LNPs can partially recapitulate the nutritional role of symbionts. Such an insight offers opportunities to experimentally isolate the metabolic versus signaling effects of symbiosis.

LNPs also represent a flexible and targeted experimental tool for probing lipid function in cnidarian holobionts. Specific lipid classes, including oxylipins, a diverse group of oxygenated fatty acid derivatives proposed to act as signaling molecules in stress responses and host–symbiont communication (Rosset et al. 2021; Gamba et al. 2024, 2025), could be selectively encapsulated into LNPs to investigate their functional roles. Delivered alone or in combination with pathway inhibitors or isotopic tracers, these compounds could be used to dissect lipid-mediated signaling under different thermal regimes, lipid transport during symbiosis establishment (particularly where lipids can be clearly attributed to either host or symbiont, e.g. by chirality), or in the presence of alternative symbiont species. Such a level of experimental precision could reveal new molecular targets involved in coral bleaching, resilience, and symbiotic compatibility.

LNP supplementation carries strong promise to support coral aquaculture, reef restoration, and climate adaptation strategies. Coral larvae supplemented with LNPs have shown enhanced survival (Matthews et al. 2025), and their use in *ex situ* propagation may offer a low-nitrogen, scalable solution to improve tissue condition and metabolic resilience in both juvenile and adult corals (Suggett and van Oppen 2022). Compared to traditional heterotrophic feeding, LNPs could reduce biofouling, eutrophication risk, and handling effort in large-scale systems. While *in situ* delivery methods remain a technical challenge, requiring innovations such as slow-release coatings, gel matrices, or targeted surface adhesion, our results suggest that even short-term exposure to LNPs is sufficient to elicit significant physiological responses. Altogether, these findings position LNPs as a potent tool in coral science: nutritionally beneficial, mechanistically insightful, and practically applicable. As climate stress continues to challenge coral reef persistence, tools like LNPs may help bridge the gap between basic mechanistic research and effective intervention.

## Data Availability

Protein mass spectrometry data are publicly available at the ProteomeXchange Consortium (Vizcaíno et al. 2014) via the PRIDE repository (Perez-Riverol et al. 2019) with identifier PXD066136.

## Author Contributions (CRediT)

E. Laurent: Investigation, formal analysis, writing – original draft preparation, writing – review and editing. J. Matthews: Conceptualization, methodology, writing – review and editing. L. Chagnat and C. de Jong: Investigation (equal), writing – review and editing. L. Peng: Resources, writing – review and editing. P. Cleves, D. Suggett: Funding acquisition (equal), writing – review and editing. C. Oakley: Conceptualization (lead), formal analysis, data curation, methodology, supervision (lead), funding acquisition (lead), visualization, writing – original draft preparation, writing – review and editing.

## Figures

**Supp. Fig. 1.**
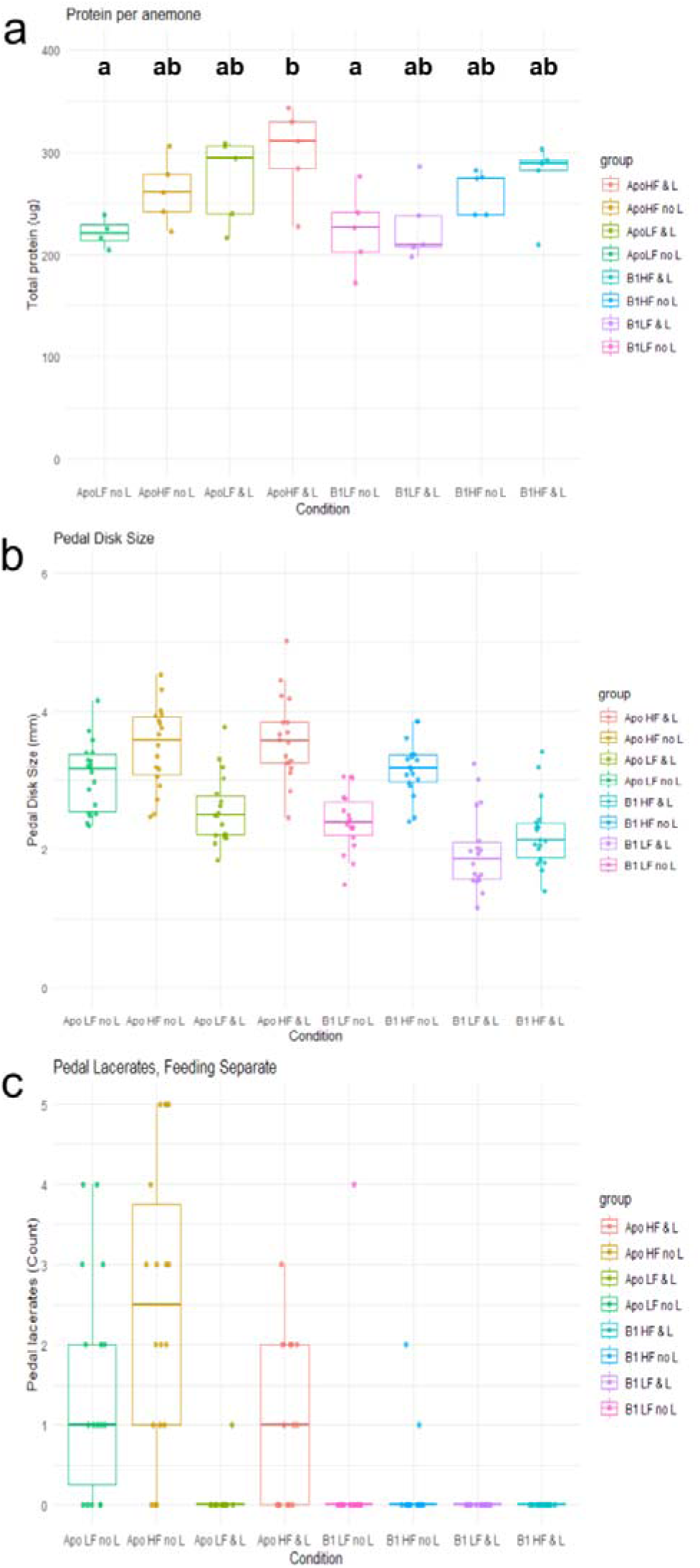
Physiological responses of *Exaiptasia diaphana* to lipid nanoparticle supplementation, with conditions separated by feeding regime. a: Protein mass per anemone at the end of the four-week experiment. b: Pedal disk size at the end of the four-week experiment. c: Total counts of pedal lacerates across the four-week experiment. Apo: aposymbiotic; B1: *Exaiptasia* hosting *Breviolum minutum*; L: lipid nanoparticle supplementation; NoL: No lipid nanoparticle supplementation, HF: high feeding (2 × week^−1^), LF: low feeding (1 × week^−1^).

